# Genome-wide association mapping for growth rate at fluctuating and extreme temperatures

**DOI:** 10.1101/2024.06.03.597084

**Authors:** Emmi Räsänen, Martta Liukkonen, Pauliina A. M. Summanen, Tarmo Ketola, Ilkka Kronholm

## Abstract

Constant temperatures and fluctuations of varying frequency affect fitness differently, which has led to suggestions of distinct genetic architectures and adaptation strategies between constant and fluctuating thermal environments. However, very little is known about the possible fitness trade-offs and genetic constraints underlying thermal adaptation and how they affect species’ ability to confront climatic changes. We addressed this gap in knowledge by integrating quantitative genetics and genome-wide association mapping in the filamentous fungus *Neurospora crassa*. Growth rates were measured for 434 strains under fast and slow frequency fluctuations, at high and low temperature range with respect to the species’ tolerance, and at constant mean and extreme temperatures of these ranges. We found strong genetic correlations between fast and slow frequency fluctuations, and between fluctuations and their mean temperatures, but not with the highest extreme temperature. Positive correlations were supported by high heritability values, pointing that in *N. crassa* there are no significant trade-offs or genetic constraints in adaptation when variance in temperature increases. Altogether, our results indicated clearly polygenic basis of thermal tolerance, with most of the variation in overall performance (83 %), and clearly less in hot-cold trade-off (8 %), or heat stress tolerance (4 %). Interestingly, GWAS discovered many SNPs associated with growth rate only at constant temperatures or at fast and slow fluctuations at high and low thermal range. However, the cellular functions of the associated genes were overlapping, and no opposite allelic effects were found between treatments. Hence, large-effect loci indicated no trade-offs, but a shared physiology across temperatures, probably owing to the general stress response or individual’s overall fitness.

## 1 INTRODUCTION

The global increases in mean and variance of temperature force species to tolerate more frequent climatic fluctuations such as severe heat waves (Meehl et al. 2000; IPCC 2018). Based on species performance projections and empirical data, adaptation to temperature fluctuations should be harder than tolerating just the rise in mean temperature (Vasseur et al. 2014; Colinet et al. 2015). Consequently, a better understanding of how species can cope with increased thermal variation is crucial for predicting their survival under climate change (Schulte et al. 2011; Kristensen et al. 2020). According to traditional hypotheses, the greater variability in temperature should lead to evolution of generalists with enhanced performance across a wider range of temperatures (Levins 1968; Lynch and Gabriel 1987; Gilchrist 1995). However, the ability to spend more time closer to critical thermal limits should evolve at the expense of overall fitness, also known as a specialist-generalist trade-off (Angilletta 2009). This hypothesis suggests that the only way to change the breadth of the thermal performance curves (TPCs) is by lowered elevation at optimal conditions, meaning that populations cannot be concurrently well-adapted to both mean and variance of temperature (Lynch and Gabriel 1987; Gilchrist 1995).

More recently, studies have acknowledged that adaptations to temperature fluctuations might not be predictable from TPCs measured across constant temperatures and shown evidence that distinct genetic architectures regulate adaptation in fluctuating versus constant conditions (Schulte et al. 2011; Ketola and Saarinen 2015; Sinclair et al. 2016; Sørensen et al. 2016; Ketola and Kristensen 2017). In addition, theoretical studies suggest that there are evolutionary constraints between the types of fluctuations, for example, that particular adaptive mechanisms are operating in environments that fluctuate at different time scales (DeWitt and Langerhans 2004; Botero et al. 2015). If these hypotheses hold true, distinct genetic architectures and constraints are associated with adaptation to constant and fluctuating environments of varying frequency (Ketola and Kristensen 2017), which could limit evolutionary adaptation and heavily affect species survival when variation in temperature increases (Sgrò and Hoffmann 2004; Björklund et al. 2009).

The potential trade-offs in thermal adaptation are often investigated by the principal components of genetic variation in TPCs and quantitative genetic analysis of covariation (Kingsolver et al. 2001; Izem and Kingsolver 2005). Specifically, evolutionary constraints are defined as directions where selection cannot increase fitness due to low amounts of heritable genetic variation or negative genetic correlations across alternative environmental states (Gaydos et al. 2013). The fluctuations of varying frequency affect fitness differently and are expected to have their characteristic changes in TPCs (Schulte et al. 2011; Colinet et al. 2015; Schaum et al. 2022). Generally, three modes of variation are observed in the shape and position of TPCs; the elevation in overall performance, horizontal shift in optimum between hot and cold temperatures, and in breadth of the TPC (Izem and Kingsolver 2005; Angilletta 2009). Under fast fluctuations, selection should favour elevated performance and more robust tolerance across temperatures, whereas slower fluctuations should select for more fixed strategies, reflected as a narrower TPCs, or shift in optimum (DeWitt and Langerhans 2004; Ketola et al. 2014; Schaum et al. 2022). However, fast fluctuations could also select for reversible phenotypic plasticity aided by the fast production of heat shock proteins or the maintenance of regulatory enzymes, which could trade-off, due to their energetic requirements with tolerance to constant environments (Kristensen et al. 2000; Sørensen et al. 2003).

The traditional quantitative genetics studies have long focused on estimating statistical averages over a multitude of loci with small phenotypic effects (Fisher 1919). More recently, the next generation sequencing methods have made it possible to use whole genome sequence data and investigate the genetic architecture of polygenic traits on the level of individual loci (Fu et al. 2013). For example, genome-wide association studies (GWAS) link genotypes to phenotypes by mapping mutational markers such as single nucleotide polymorphisms (SNPs). Using quantitative genetics and the latest molecular genetics methods together can give us more complete picture of the genetic variation in thermal tolerance and its evolutionary potential (Cortés et al. 2020; Buckley and Kingsolver 2021), but so far only few studies have adopted this approach (Gerken et al. 2015; Latimer et al. 2015; Duun Rohde et al. 2016; Rolandi et al. 2018; Lecheta et al. 2020). Furthermore, most of the quantitative genetics studies and studies using genome-wide sequence data have focused on evaluating the effects of constant or increasing heat (Riehle et al. 2001; Knies et al. 2006; Mitchell and Hoffmann 2010; Khan et al. 2022). Knowledge from differently fluctuating temperatures is sparse and studies have mostly used experimental evolution and resequencing (Tobler et al. 2014; Deatherage et al. 2017; Schaum et al. 2018; Lambros et al. 2021), whereas only one study has been made on natural genetic variation (Sørensen et al. 2016). As climate change keeps accelerating, there is a pressing need for experiments evaluating the predicted fitness trade-offs and constraints in thermal adaptation under various fluctuations (Chevin and Hoffmann 2017; Buckley and Kingsolver 2021).

To address this gap in knowledge, we investigated if distinct genetic architectures are found at constant and fluctuating temperatures of varying frequency, and if some genetic constraints might limit thermal adaptation. We used quantitative genetics and GWAS in the filamentous fungus *Neurospora crassa* to quantify how different components of genetic variation contribute to the total variation. The thermal sensitivity to growth rate was measured for a previously developed mapping population (Moghadam et al. 2020; Räsänen et al. 2024) under fast and slow frequency fluctuations, at high and low temperature range, and at constant mean and extreme temperatures of these ranges. In our analyses we used a multivariate approach in which growth rates at each temperature are treated as discrete and potentially genetically correlated traits (Kingsolver et al. 2004). We examined the following questions: (1) How much there is genetic variation in growth rate at constant and fluctuating temperatures and how strongly is this variation correlated between contrasting thermal environments? (2) To what extent the phenotypic variation in growth rate can be related to broader characteristics of the TPCs? (3) Are there genes associated with growth rate at constant and fluctuating temperatures and are these genes same between contrasting thermal environments? Our study is among the first ones to investigate the genetic basis of thermal tolerance at temperatures that fluctuate with different frequencies.

## 2 MATERIALS AND METHODS

### 2.1 *Neurospora crassa* mapping population

*N. crassa* is a clonally and sexually reproducing species that is well-suited for experiments studying genetic differences across environments (Fisher and Lang 2016). In our experiment we used 118 natural strains obtained from the Fungal Genetics Stock Center (Manhattan, Kansas, USA) (McCluskey et al. 2010), originally sampled from natural populations in southeastern USA (Louisiana), Central America and the Caribbean Basin (Ellison et al. 2011; Palma-Guerrero et al. 2013). Previously, some of the natural strains were crossed to form 316 additional offspring strains (Moghadam et al. 2020). In total our nested association mapping population had 434 strains genotyped for 1 473 869 single nucleotide polymorphisms (SNPs) (Räsänen et al. 2024).

### 2.2 Phenotyping by growth rate

We measured the mycelial growth rate as a proxy for thermal tolerance (Pringle and Taylor 2002). Before starting experiments, *N. crassa* cultures were maintained on Vogel’s medium N (Metzenberg 2003) in slants at room temperature. The linear growth rates were measured for each strain by standard tube method using agar filled serological pipet tips (Ryan et al. 1943; White and Woodward 1995). A more detailed description of this method can be found from Moghadam et al. (2020) and Kronholm et al. (2016). We replicated each strain 3 times and put them to grow in 10 thermal treatments (MTM-313 Plant Growth Chamber, HiPoint Corp., Taiwan). We had two fluctuating regimes, low 25–35 °C and high 32–42 °C, and fast and slow fluctuations with frequencies of 120 min and 480 min respectively. Thus, fast fluctuations were 4 times faster, having a step duration of 15 min at each temperature compared to 60 min steps at slow fluctuations. At lower range the steps were 25–27.5–30–32.5–35–32.5–30–27.5–25 °C, and at higher range 32–34.5–37–39.5–42–39.5–37–34.5–32 °C. The fast fluctuations were estimated to span within the generation time of *N. crassa*, whereas the slow fluctuations occurred between generations (Kronholm and Ketola 2018). The growth rates were measured also at constant mean and extreme temperatures of these ranges. The measurements at constant lower range temperatures 25 °C, 30 °C, and optimal 35 °C were collated from previously published data (Moghadam et al. 2020). At higher range the growth rates were measured at constant 32 °C, 37 °C, and extreme 42 °C which is known to be very stressful for *N.* crassa (Mohsenzadeh et al. 1998). Altogether we had 12 976 growth assays to use in our analyses.

### 2.3 Quantitative genetics

The statistical analysis for quantitative genetics parameters was conducted with Bayesian multilevel model using Hamiltonian Monte Carlo algorithm. We used R version 4.3.1 (R Core Team 2022) and R package ‘brms’ version 2.19.0 (Bürkner 2018) that fits Bayesian models by ‘Stan’ language. The phenotypic data including all growth assays was used in a multivariate model

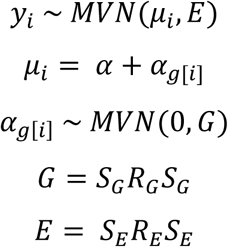

where the growth at each temperature treatment was explained by the strain genotype as a random variable and modeled as potentially correlated with growth at other temperatures. In this model α was the vector of intercepts, *α_g[i]_* was the vector of genotypic effects, *S_G_* and *S_E_* were 10 × 10 diagonal matrices with genetic or environmental standard deviations on the diagonal, and *R_G_* and *R_E_* were matrices for genetic and environmental correlations, respectively. We had weakly informative priors as we used the half location-scale version of Student’s t-distribution with 3 degrees of freedom and 2.5 as the scale parameter. The prior for the intercept effects was

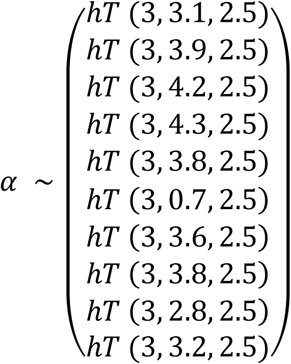

for growth rates from 25 °C to 42 °C, and for fluctuating temperatures in order 25–35 °C fast, 25–35 °C slow, 32–42 °C fast, and 32–42 °C slow. The prior for each standard deviation in the model was *σ* ∼ hT(3, 0, 2.5), and we used an lkj prior (McElreath 2015) for the correlation matrices: *R_E_*, *R_G_* ∼ LKJ(1). For MCMC estimation 2 chains were run with a warmup period of 1000 iterations, followed by 6000 iterations of sampling, with thinning set to 2. MCMC convergence was monitored by trace plots and 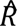 values.

From the model we obtained a G-matrix with estimates of genetic variance and covariance components for growth rate. The environmental variances were estimated as the variances between the clonal replicates of the same strain. We considered the posterior mean estimates to be statistically significant if their 95 % highest posterior density intervals (HPDI) did not overlap with 0. We extracted the posterior samples from the model and used these to calculate broad-sense heritabilities, genetic covariances and genetic correlations for growth rate in different thermal environments. We also quantified the coefficients of variation for both, the genetic and environmental components by standardizing the scales by the mean phenotype. The R package ’ggplot2’ (Wickham 2016) was used for plotting.

### 2.4 Principal components analysis

We used a Principal Components Analysis (PCA) to characterize the main modes of variation in TPCs, and the amount of phenotypic variation that was attributed to each principal component (PC) (Izem and Kingsolver 2005). We implemented PCA with built-in R function ‘princomp’ for the growth rates averaged per temperature for each strain. The predictive values of the first 3 principal components (from here on PC1, PC2 and PC3) were saved and used in GWAS so that the results could be compared with the associations of specific temperature treatments. We counted also the Bayesian mean estimates and HPDIs for the PCs. To form the posterior distributions, we sampled the intercepts extracted from the multivariate model for each genotype and at each temperature 5000 times. Then we used the PCA to calculate the proportions of variance explained by the first 6 PCs and the loadings of PC1, PC2 and PC3 at each temperature.

### 2.5 Genome-wide association mapping

GWAS was used to study the genotype–phenotype associations and to answer whether the same or different genes contribute to the thermal tolerance at fluctuating and constant temperatures. We investigated also which SNPs are associated with PC1, PC2 and PC3 explaining the variation in average growth rate. We used nested association mapping that combines information from recent and past recombination (Yu et al. 2008). We conducted the GWAS by the genome association and prediction integrated tool (GAPIT) Version 3 R-package (Wang and Zhang 2021). As an input data we had each strains SNP sequence in HapMap format, and the strains average growth rate per temperature treatment. In GWAS, GAPIT 3 identifies the quantitative trait loci via statistical association between the phenotypes and single-nucleotide markers. It also corrects for the population structure and kinship in these analyses and so sets control for the false positives. The basic mixed model was

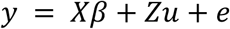

where *y* is a vector of phenotypic observations, *β* is a vector of fixed effects including effects of the genetic marker and population structure, *u* is a vector of random genetic effects for each individual, X and Z are design matrices, and *e* is a vector of residual effects. The variance matrix for *u* is 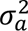 K where 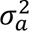 is the additive variance and K is the kinship matrix. The variance matrix for *e* is 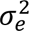 I, where 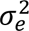 is the residual variance and I is the identity matrix. For the population structure correction, a principal component analysis was first done on the SNPs, and the components were incorporated as fixed factors in the model (Price et al. 2006). We chose the first 4 components for the analysis since they explained most of the variance in SNPs.

We run GWAS with Bayesian-information and linkage-disequilibrium iteratively nested keyway method (BLINK) (Huang et al. 2019). BLINK is a multi-locus test (Segura et al. 2012) and the most statistically powerful method in GAPIT 3 (Wang and Zhang 2021). In BLINK, the SNP markers are removed iteratively if they are in linkage disequilibrium with the most significantly associated reference SNP. The procedure of selecting a reference SNP among the remaining markers is repeated until no markers can be removed. We run the BLINK separately for the average growth rates in all temperature treatments and for the phenotypic variation attributed to PC1, PC2 and PC3 describing the TPC. The significance threshold for associations was set to 0.01 after a Bonferroni correction, which corresponded to a P-value of 6.78 × 10^−9^. The signs of the allelic effects were with respect to the minor allele, i.e., a positive allelic effect indicated that the individuals carrying the minor allele had faster growth rate.

We searched the statistically significant SNPs from Ensemble Fungi release 57 (Yates et al. 2022) and identified the candidate genes affecting the growth rate. We used *Neurospora crassa* (NC12) whole genome alignment and annotations (Galagan et al. 2003). We also processed the SNP variants by the Variant Effect Predictor web tool (McLaren et al. 2016) to determine the possible effects on transcript and protein sequences. In the case of intergenic variants, we reported annotations for a gene that had the shortest distance from the marker to the transcript. We further searched for the functional protein annotations of the candidate genes from the UniProt (The UniProt Consortium 2023) and the FungiDB databases (Basenko et al. 2018).

## 3 RESULTS

### 3.1 Growth rates at different temperatures

We observed large variation in growth rates between temperature treatments (0.01– 6.05mm/h) and there were clear differences in how well strains grow in fluctuating temperatures (Fig. 1A). The genotype had a statistically significant effect on growth rate at each temperature treatment (the model estimates are reported as a mean with the 95 % HPD interval given in parentheses). On average, the growth rates were faster at lower temperature range and at slow frequency fluctuations (Fig. 1B). At constant temperatures, growth rates increased from 25 °C (3.04 mm/h, 3.00–3.08) to 30 °C (3.83 mm/h, 3.77–3.88), and at 32 °C (4.07 mm/h, 4.01–4.12). The growth rates were fastest at the optimal temperature 35 °C (4.16 mm/h, 4.10–4.23) and slowed down at hotter 37 °C (3.73 mm/h, 3.66–3.79) and at 42 °C (0.68 mm/h, 0.65–0.72). At lower 25–35 °C range, the mean growth rates were very similar, 3.53 mm/h (3.48–3.58) at fast fluctuations and 3.65 mm/h (3.61–3.70) at slow fluctuations. At higher 32–42 °C range, the mean growth rate was faster at slow frequency fluctuations 3.10 mm/h (3.05–3.15) than at fast fluctuations 2.76 mm/h (2.72–2.81).

**Figure 1.**
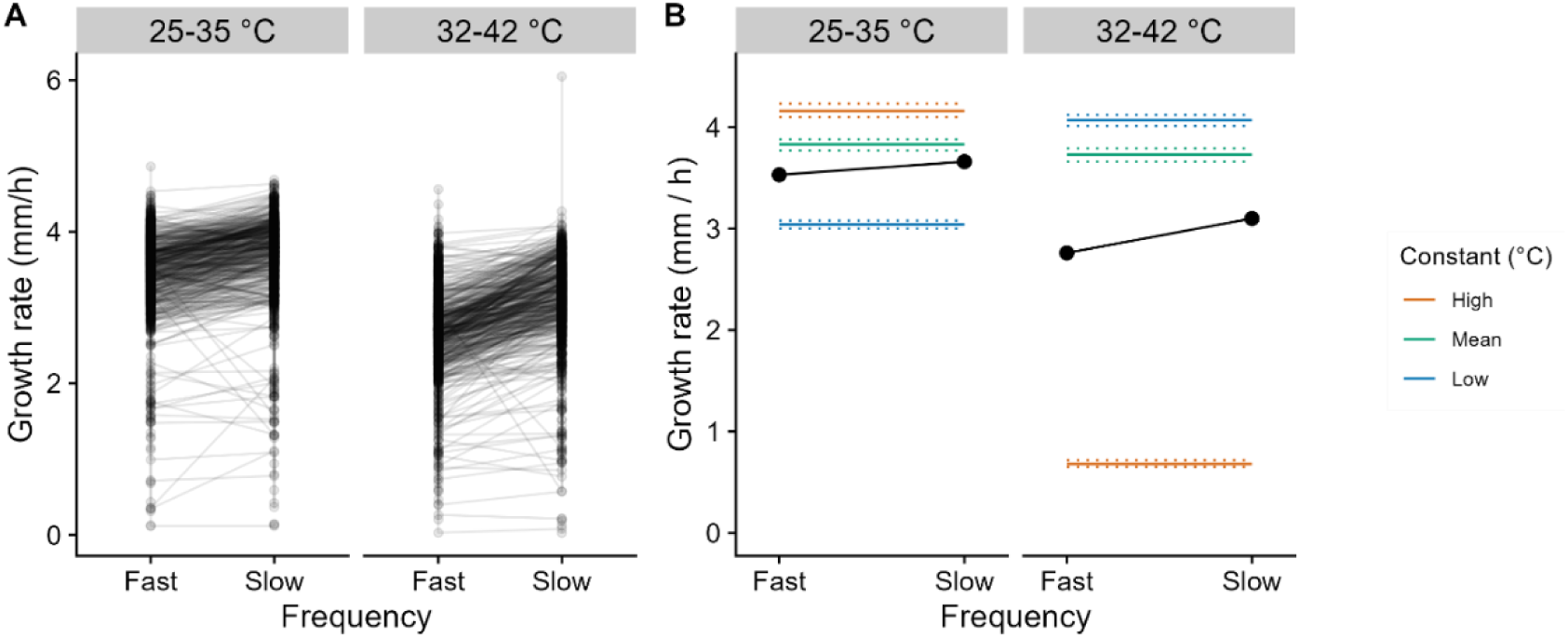
(A) The mean growth rates for each *N. crassa* genotype (N = 434) and (B) the posterior mean estimates of growth rate when the growth in fluctuating environments was compared at high and low temperature range and at fast and slow frequency. In Fig. B the estimates for the constant temperatures within a temperature range are indicated by colors (red = high extreme temperature, green = mean temperature, and blue = low extreme temperature, reflecting 25 °C, 30 °C, and 35 °C at lower range, and 32 °C, 37 °C, and 42 °C at higher range). The 95 % HPD intervals are marked with dotted lines for constant temperatures. In fluctuating temperatures, the points are hiding the small 95 % HPD interval bars.

### 3.2 Quantitative genetics

The multivariate model was used to estimate the genetic variance and covariance components for growth rate at different temperatures (Table 1). In all thermal treatments, there was genetic variation for growth rate (posterior means 0.14–0.46) (Table 1), and this was observable also from high heritability values (posterior means 0.64–0.92) (Fig. 2A). In general, we found strong positive genetic correlations (posterior means 0.54–0.99) and genetic covariances (posterior means 0.10–0.35) between treatments (Table 1). However, the correlations with constant 42 °C were markedly lower (posterior means 0.14–0.39), which was also true for the covariances (posterior means 0.02–0.07). There were most genetic and environmental variance at constant 42 °C when scaled as the coefficients of variation (Fig. 2B).

**Table 1.**
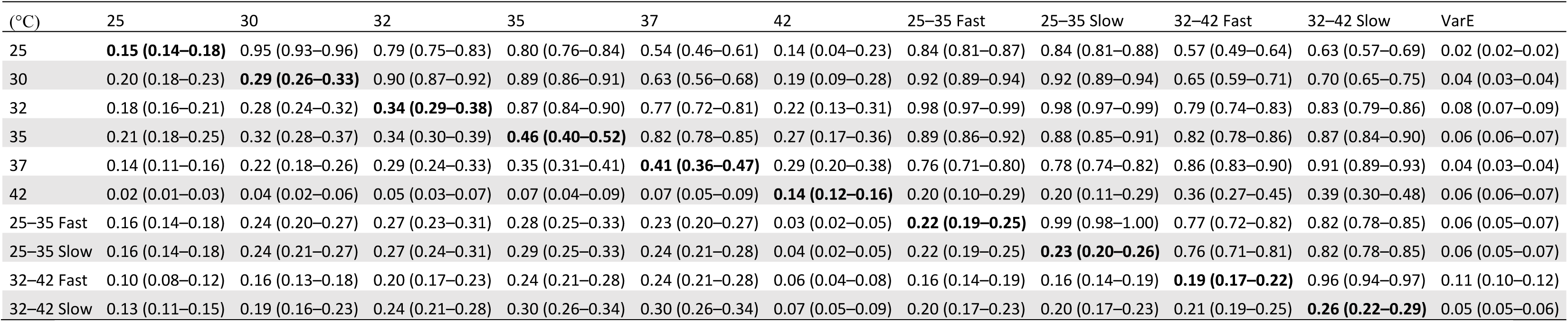
Genetic variances, covariances, correlations, and environmental variances for growth rate at constant and fluctuating temperatures estimated from the multivariate model. In table: upper triangle genetic correlations, lower triangle genetic covariances, diagonal genetic variances (in bold), and last column environmental variances. Estimates are posterior means with 95 % HPD intervals shown in parentheses.

**Figure 2.**
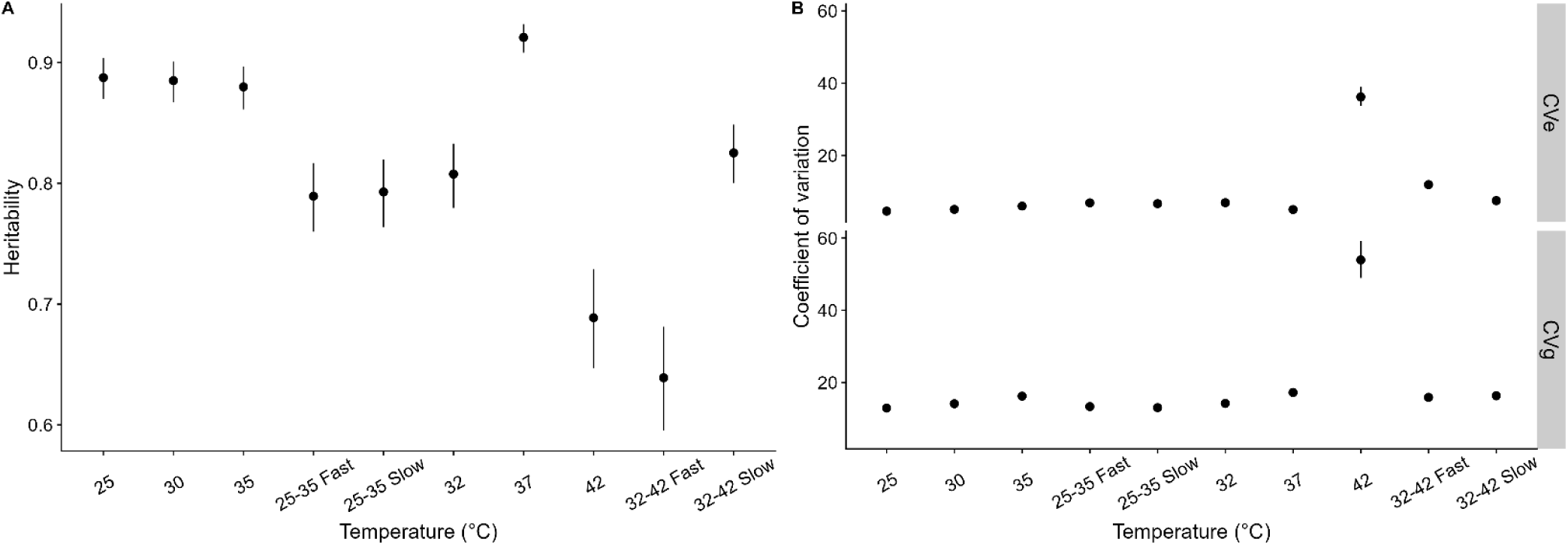
(A) Heritabilities of growth rate and (B) coefficients of genetic and environmental variation at each temperature treatment. Estimates are posterior means with 95 % HPD intervals (some points are larger than the error bars).

### 3.3 Principal component analysis

PCA found 3 principal components that explained 95 % of the total genetic variance in average growth rate (Fig. 3A). The loadings of these 3 principal components at different thermal treatments were easily interpreted as TPC attributes (Fig. 3B). PC1 explained 83 % of the total variation and had the largest loading (0.44) at the optimal temperature 35 °C. As all loadings were positive, PC1 was interpreted as the individual’s overall fitness in growth i.e. the elevation of the TPC (Fig. 3C). PC2 explained 8 % of the variation and was interpreted as the horizontal shift in TPC optimum towards colder temperatures (Fig. 3D). Interestingly, the constant 37 °C and 42 °C, and 32–42 °C fluctuations all had negative loadings in PC2. The proportion of variance for PC3 was only 4 % and the loadings were interpreted as the change in the shape of the TPC (Fig. 3E). There was a strong positive loading (0.90) for PC3 at constant 42 °C, indicating increased growth rate at the high thermal extreme.

**Figure 3.**
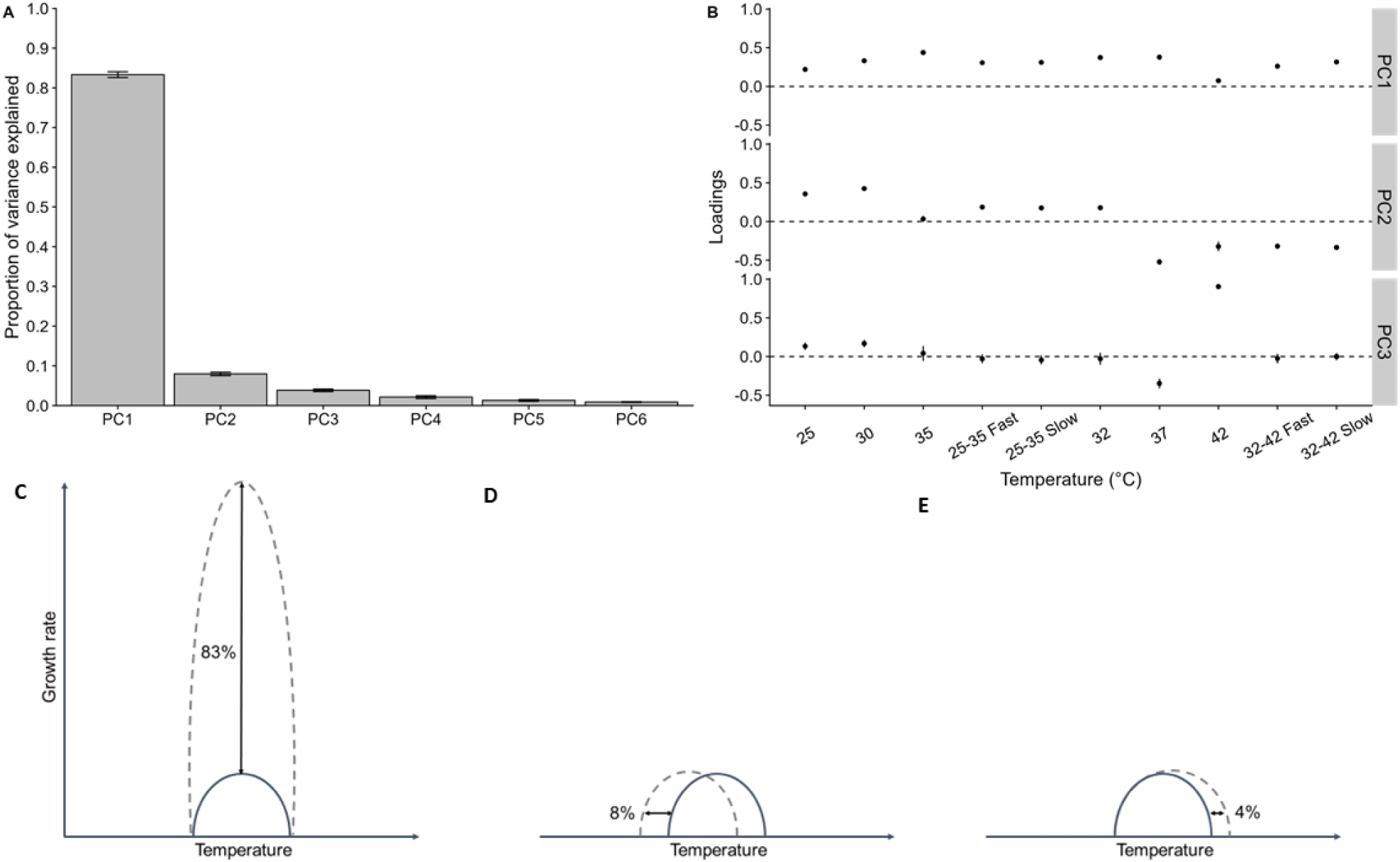
(A) Proportions of variance explained by the different principal components and (B) the loadings of PC1, PC2 and PC3 for each temperature treatment. Shown are the posterior means and 95 % HPD intervals as error bars. At lower line are the schematic pictures for the changes in TPC attributes: (C) the elevation, (D) the hot and cold shift in optimum, and (E) the tolerance of high thermal extreme.

### 3.4 Genome-wide association mapping

#### 3.4.1 Associations and allelic effects

GWAS found 25 associations with growth rate at different temperatures, which were related to 21 SNPs, and located in 19 genes or intergenic loci (Table 2). At constant temperatures optimal or lower, significant SNPs were found from chromosomes 3, 5 and 6 (Fig. 4, Fig. 5). At constant temperatures above the optimal, the associated SNPs were in chromosomes 1 and 6 (Fig. 5). At fluctuations, associations were in chromosomes 1, 3 and 6 (Fig. 4, Fig. 5). The strongest associations were found at 42 °C in chromosome 1: 7 807 362 (P = 4.66 × 10^−32^), at slow 25–35 °C fluctuations in 1: 7 378 135 (P = 2.47 × 10^−21^), and at fast 25–35 °C fluctuations in 3: 2 881 603 (P = 2.77 × 10^−19^) and in 6: 3 600 909 (P = 1.19 × 10^−19^). From significantly associated SNPs, 8 were detected only at fluctuating temperatures, 12 only at constant temperatures, and 2 at both, fluctuating and constant temperatures. Altogether there were 18 unique SNPs associated only at one temperature treatment and 3 shared SNPs that were associated at more than one temperature treatment (Table 2). Only unique SNPs were found at the lowest extreme temperature 25 °C, above optimum at 37 °C and 42 °C, and at higher 32–42 °C range. The shared SNPs 3: 2 300 888, 3: 2 881 603, and 6: 3 600 716 were found at favorable 30 °C, 32 °C, 35 °C, and fluctuating 25–35 °C temperatures. Interestingly, there were no shared SNPs between the fast and slow fluctuations.

**Table 2.**
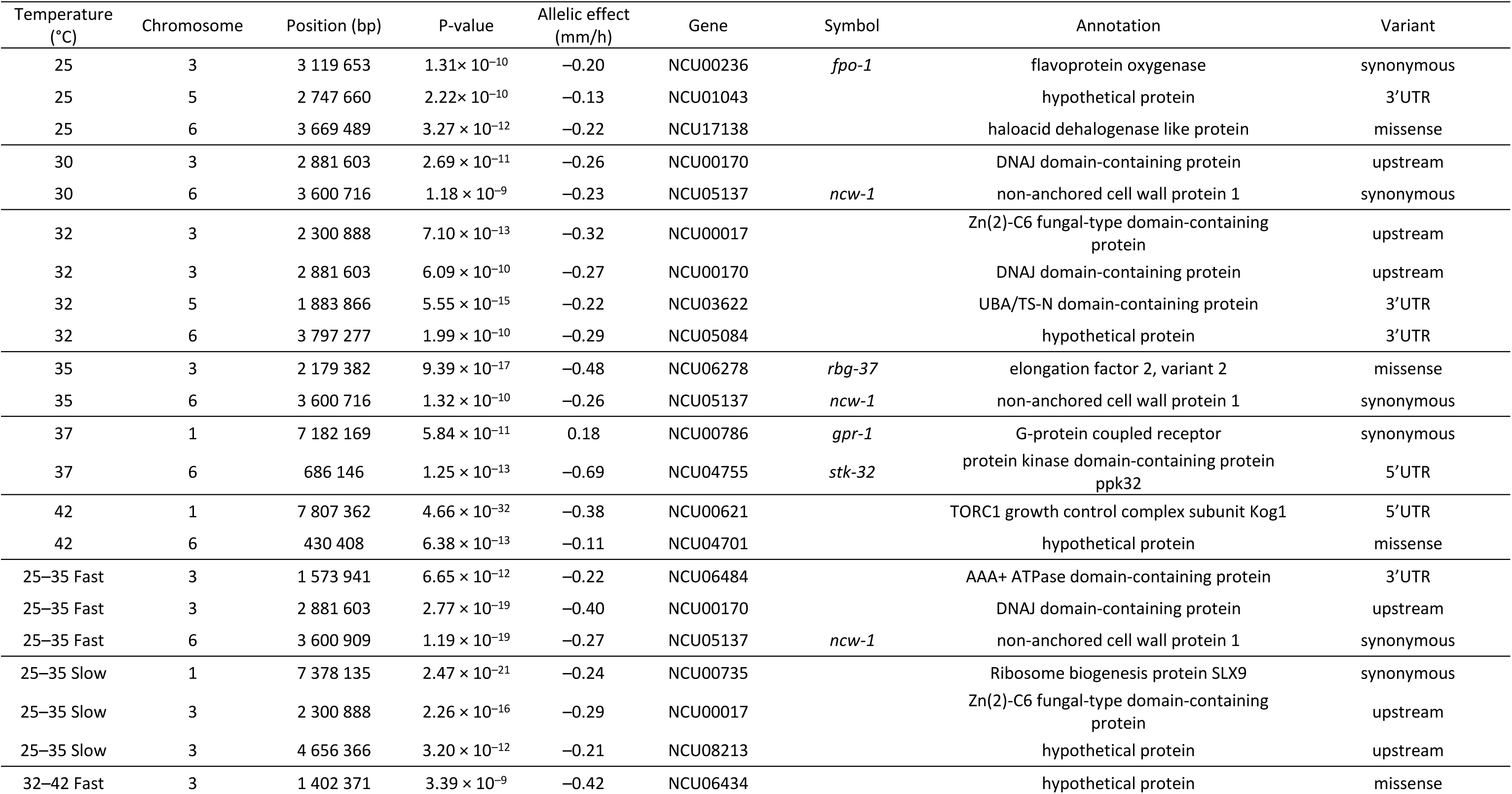
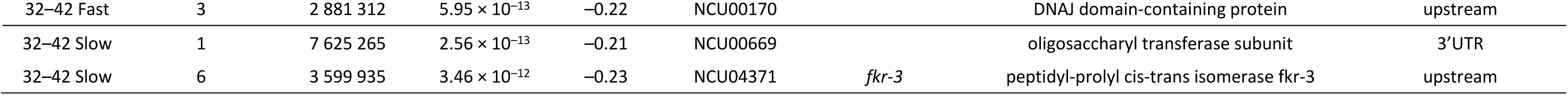
SNPs that were significantly associated with growth rate at constant and fluctuating temperatures, their positions in genome (bp), allelic effects (mm/h), variant types and annotated proteins. The signs of the allelic effects are with respect to the minor allele and annotations for the intergenic variants are reported for the closest gene.

| Temperature (°C) | Chromosome | Position (bp) | P-value | Allelic effect (mm/h) | Gene | Symbol | Annotation | Variant |
| --- | --- | --- | --- | --- | --- | --- | --- | --- |
| 25 | 3 | 3 119 653 | $1.31 \times 10^{-10}$ | -0.20 | NCU00236 | *fpo-1* | flavoprotein oxygenase | synonymous |
| 25 | 5 | 2 747 660 | $2.22 \times 10^{-10}$ | -0.13 | NCU01043 | | hypothetical protein | 3'UTR |
| 25 | 6 | 3 669 489 | $3.27 \times 10^{-12}$ | -0.22 | NCU17138 | | haloacid dehalogenase like protein | missense |
| 30 | 3 | 2 881 603 | $2.69 \times 10^{-11}$ | -0.26 | NCU00170 | | DNAJ domain-containing protein | upstream |
| 30 | 6 | 3 600 716 | $1.18 \times 10^{-9}$ | -0.23 | NCU05137 | *ncw-1* | non-anchored cell wall protein 1 | synonymous |
| 32 | 3 | 2 300 888 | $7.10 \times 10^{-13}$ | -0.32 | NCU00017 | | Zn(2)-C6 fungal-type domain-containing protein | upstream |
| 32 | 3 | 2 881 603 | $6.09 \times 10^{-10}$ | -0.27 | NCU00170 | | DNAJ domain-containing protein | upstream |
| 32 | 5 | 1 883 866 | $5.55 \times 10^{-15}$ | -0.22 | NCU03622 | | UBA/TS-N domain-containing protein | 3'UTR |
| 32 | 6 | 3 797 277 | $1.99 \times 10^{-10}$ | -0.29 | NCU05084 | | hypothetical protein | 3'UTR |
| 35 | 3 | 2 179 382 | $9.39 \times 10^{-17}$ | -0.48 | NCU06278 | *rbg-37* | elongation factor 2, variant 2 | missense |
| 35 | 6 | 3 600 716 | $1.32 \times 10^{-10}$ | -0.26 | NCU05137 | *ncw-1* | non-anchored cell wall protein 1 | synonymous |
| 37 | 1 | 7 182 169 | $5.84 \times 10^{-11}$ | 0.18 | NCU00786 | *gpr-1* | G-protein coupled receptor | synonymous |
| 37 | 6 | 686 146 | $1.25 \times 10^{-13}$ | -0.69 | NCU04755 | *stk-32* | protein kinase domain-containing protein ppk32 | 5'UTR |
| 42 | 1 | 7 807 362 | $4.66 \times 10^{-32}$ | -0.38 | NCU00621 | | TORC1 growth control complex subunit Kog1 | 5'UTR |
| 42 | 6 | 430 408 | $6.38 \times 10^{-13}$ | -0.11 | NCU04701 | | hypothetical protein | missense |
| 25–35 Fast | 3 | 1 573 941 | $6.65 \times 10^{-12}$ | -0.22 | NCU06484 | | AAA+ ATPase domain-containing protein | 3'UTR |
| 25–35 Fast | 3 | 2 881 603 | $2.77 \times 10^{-19}$ | -0.40 | NCU00170 | | DNAJ domain-containing protein | upstream |
| 25–35 Fast | 6 | 3 600 909 | $1.19 \times 10^{-19}$ | -0.27 | NCU05137 | *ncw-1* | non-anchored cell wall protein 1 | synonymous |
| 25–35 Slow | 1 | 7 378 135 | $2.47 \times 10^{-21}$ | -0.24 | NCU00735 | | Ribosome biogenesis protein SLX9 | synonymous |
| 25–35 Slow | 3 | 2 300 888 | $2.26 \times 10^{-16}$ | -0.29 | NCU00017 | | Zn(2)-C6 fungal-type domain-containing protein | upstream |
| 25–35 Slow | 3 | 4 656 366 | $3.20 \times 10^{-12}$ | -0.21 | NCU08213 | | hypothetical protein | upstream |
| 32–42 Fast | 3 | 1 402 371 | $3.39 \times 10^{-9}$ | -0.42 | NCU06434 | | hypothetical protein | missense |
| 32–42 Fast | 3 | 2 881 312 | $5.95 \times 10^{-13}$ | −0.22 | NCU00170 | | DNAJ domain-containing protein | upstream |
| 32–42 Slow | 1 | 7 625 265 | $2.56 \times 10^{-13}$ | −0.21 | NCU00669 | | oligosaccharyl transferase subunit | 3'UTR |
| 32–42 Slow | 6 | 3 599 935 | $3.46 \times 10^{-12}$ | −0.23 | NCU04371 | *fkr-3* | peptidyl-prolyl cis-trans isomerase fkr-3 | upstream |

**Figure 4.**
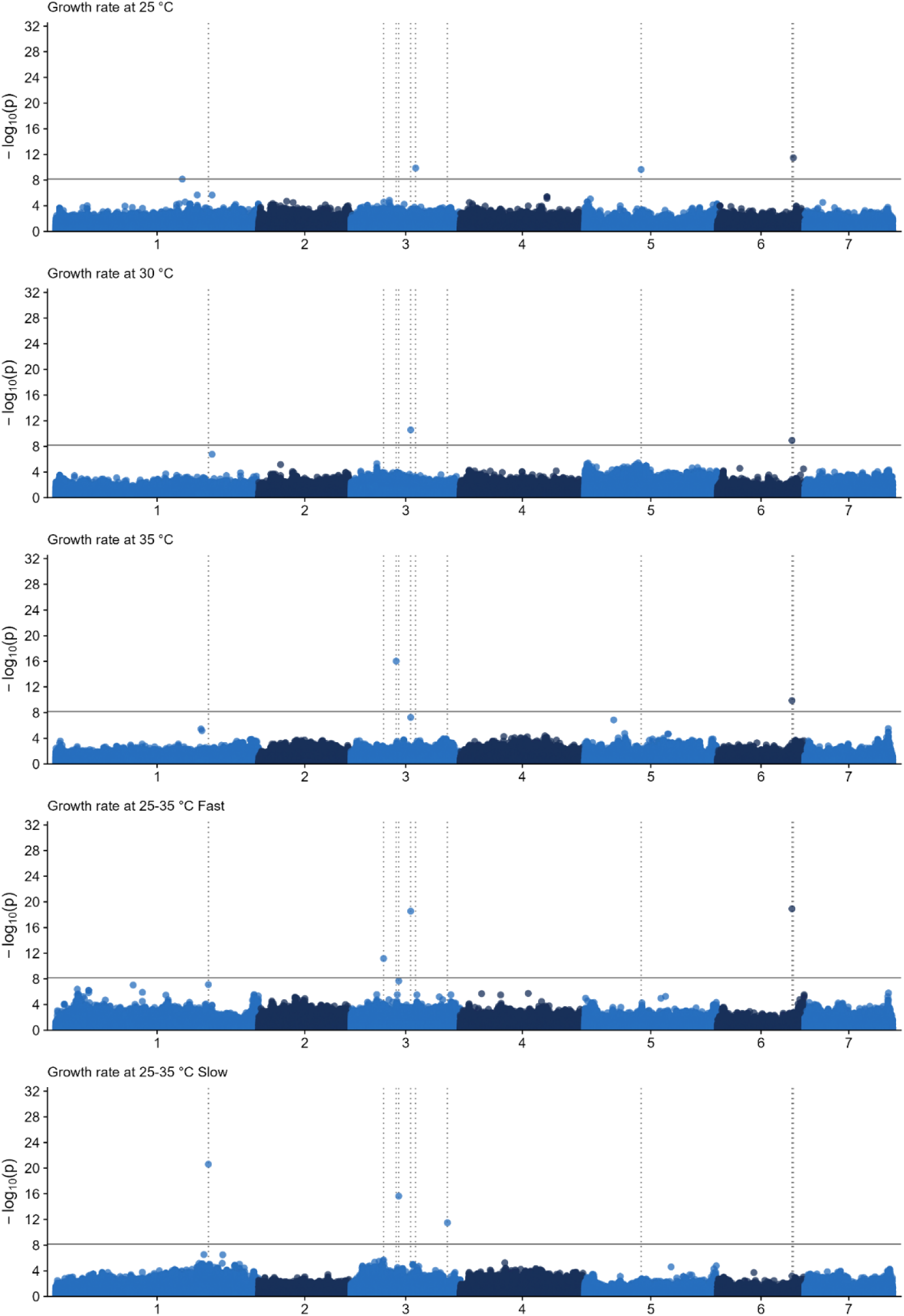
Manhattan plots for the growth rates at different temperatures. The solid horizontal lines denote a P-value threshold of 0.01 after a Bonferroni correction. The dotted vertical lines highlight the candidate SNPs that were significant in at least one of the temperature treatments within the lower range 25–35°C.

**Figure 5.**
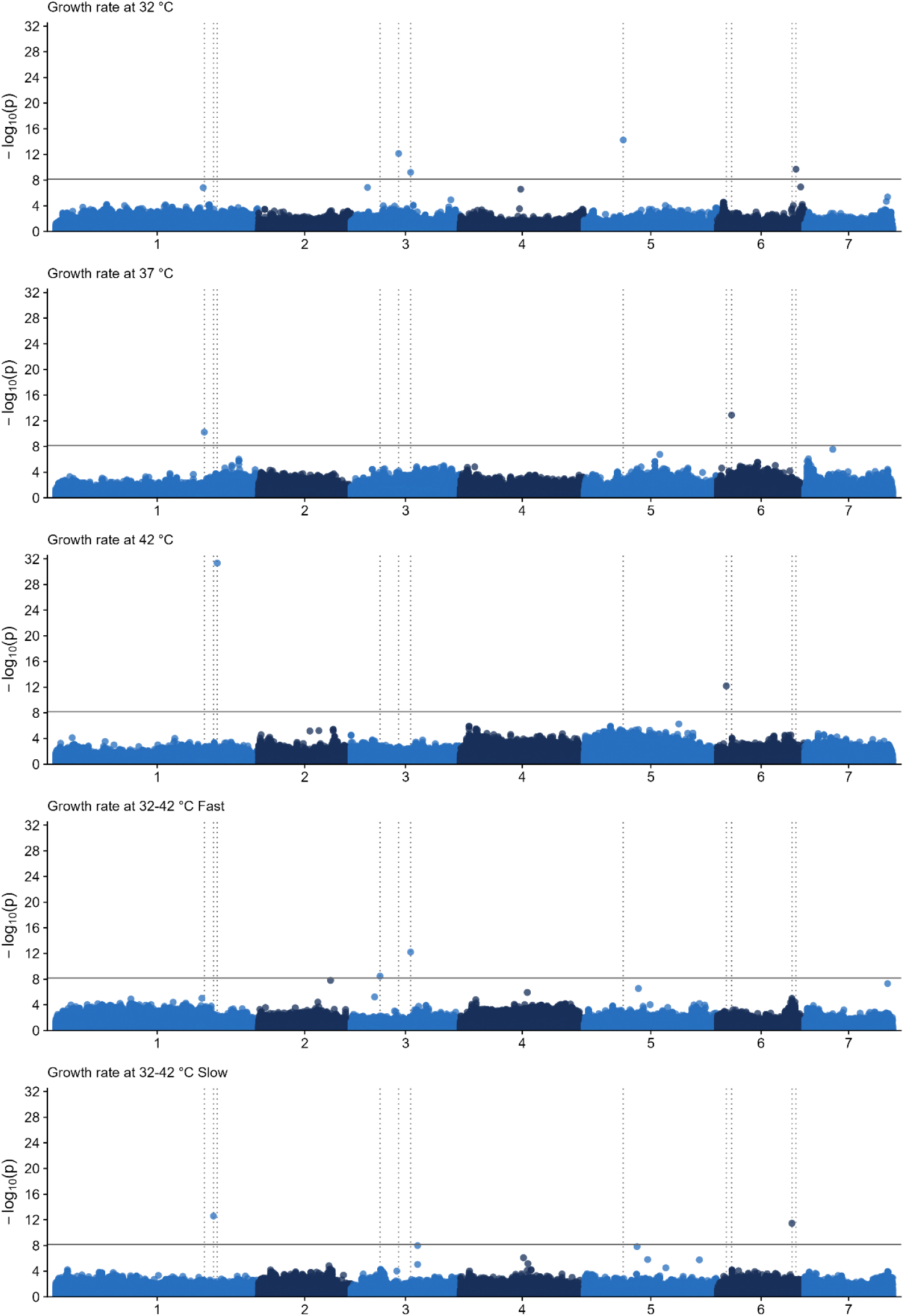
Manhattan plots for the growth rates at different temperatures. The solid horizontal lines denote a P-value threshold of 0.01 after a Bonferroni correction. The dotted vertical lines highlight the candidate SNPs that were significant in at least one of the temperature treatments within the higher range 32–42°C.

Most of the SNPs had a negative effect on growth rate when examined with respect to the minor allele (Table 2). This was also true for the shared SNPs, for which the direction of the allelic effects on growth rate were negative regardless of the temperature. The only positive effect on growth rate (0.18 mm/h) was found at 37 °C for the SNP 1: 7 182 169. In general, the minor alleles in our natural populations of *N. crassa* are known to be mildly deleterious and occur at low frequencies (Räsänen et al. 2024). The magnitude of the allelic effects was, on average, similar for the unique and the shared SNPs. The greatest allelic effects were found at 37 °C (–0.69 mm/h), at 35 °C (–0.48 mm/h), and at fast 25–35 °C fluctuations (–0.42 mm/h).

#### 3.4.2 Allelic variants

Most of the allelic variants were synonymous or in non-coding area up- or downstream from the gene (Table 2). GWAS found also 4 missense variants changing the amino acid and affecting the protein structure. These were in chromosome 6: 3 669 489 at 25 °C changing serine to proline, in 3: 2 179 382 at 35 °C changing glycine to serine, in 6: 430 408 at 42 °C changing serine to phenylalanine, and in 3: 1 402 371 at fast 32–42 °C fluctuations, changing serine to glycine. All missense variants were unique associations, whereas shared associations were synonymous or in non-coding area. However, there was no obvious connection between the types of allelic variants and the associations being found at constant or fluctuating temperatures.

#### 3.4.3 Gene annotations and protein functions

The associations found at constant lower range temperatures 25 °C, 30 °C, and at optimal 35 °C have been previously annotated in Räsänen et al. (2024). At constant 25 °C were associated the *fpo-1* (NCU00236) gene which encodes flavoprotein oxygenase, a gene (NCU01043) coding an uncharacterized protein, and a missense mutation in the gene (NCU17138) encoding a haloacid dehalogenase (HAD) like protein (Table 2). At constant 30 °C and 32 °C, and at fast fluctuations of both higher and lower range, 2 SNPs were associated in the regulatory area of the gene (NCU00170) encoding DNAJ domain-containing protein, which belongs to the HSP40 heat shock proteins. At constant 30 °C and 35 °C, and at fast 25–35 °C fluctuations there were 2 synonymous mutations in the *ncw-1* (NCU05137) gene encoding a highly conserved non-anchored cell wall protein 1. At constant 32 °C were associated a gene (NCU05084) encoding an uncharacterized protein, and a gene (NCU03622) encoding a UBA/TS-N domain-containing protein. This protein is known to function in endocytosis and to have temperature-dependent regulation of expression in fungus *Aspergillus flavus* (Georgianna et al. 2008). At constant 32 °C and at slow 25–35 °C fluctuations was associated a gene (NCU00017) encoding a Zn(2)-C6 fungal-type domain-containing protein, that is a transcription regulator. At optimal 35 °C was found a unique missense SNP in the *rbg-37* (NCU06278) gene producing a variant in elongation factor 2.

At constant 37 °C was found a synonymous association in the *gpr-1* (NCU00786) gene encoding a G-protein coupled receptor. The G-protein receptors work on the cell surface and affect the gene regulation through signaling pathways. The g*rp-1* gene has been found to be important in the sexual reproduction of *N. crassa* (Krystofova and Borkovich 2006), and the deletion mutants of another G-protein coupled receptor gene (*grp-4*), have been shown to grow well at high temperatures up to 42 °C (Li and Borkovich 2006). At constant 37 °C was also associated the *stk-32* (NCU04755) gene encoding a protein kinase domain-containing protein ppk32 (large allelic effect of –0.69 mm/h). The protein kinases are universally important coenzymes and enzyme regulators that bind to ATP in phosphorylation and work in signal transduction. In the previous GWAS, the association with the *stk-32* gene was found for the maximum growth rate at the optimal temperature (Räsänen et al. 2024).

At extreme 42 °C we found an association with the gene (NCU00621) encoding a TORC1 growth control complex subunit Kog1, and a missense mutation in the hypothetical protein (NCU04701) that had a high minor allele frequency in the mapping population (0.47) (Table 2). The target of rapamycin complex 1 (TORC1) is a highly conserved serine/threonine kinase that regulates fungal mycelial growth and protein synthesis in response to nutrient and energy signaling pathways (Wang et al. 2023). In yeast cells, TORC1 has been shown to regulate growth also by sensing environmental stressors such as high temperature (Urban et al. 2007). In *N. crassa*, several studies have linked the TOR signaling pathway to cell cycle and circadian rhythmicity, which are also affected by the changes in temperature (Lakin-Thomas 2019).

At fast 25–35 °C fluctuations was found an unique association in the gene (NCU06484) encoding a AAA+ ATPase domain-containing protein (Table 2), that positively regulates transcription. At slow 25–35 °C, the found SNPs were in the gene (NCU00735) encoding a Ribosome biogenesis protein SLX9, and upstream from the gene (NCU08213) encoding a hypothetical protein. At fast 32–42 °C fluctuations was associated a missense mutation in the gene (NCU06434) encoding a hypothetical protein. At slow 32–42 °C fluctuations, the first association was in the gene (NCU00669) producing an oligosaccharyl transferase (OST) subunit. OST is a membrane protein complex, which catalyzes a central step in the glycosylation pathway. Glycosylation affects the folding and thermodynamic stability of the proteins. The N-glycosylation pathway has been found to be highly conserved across species of filamentous fungi (Deshpande et al. 2008). The second association was upstream from the *fkr-3* (NCU04371) gene which produces a peptidyl-prolyl cis-trans isomerase. The *fkr-3* gene accelerates the protein folding by catalyzing the cis-trans isomerization activity. In *N. crassa*, the *fkr-3* deletion mutants have a temperature-sensitive phenotype (Pinto et al. 2008), and this gene is upregulated during heat stress and recovery also in other fungi (Zou et al. 2018).

#### 3.4.4 GWAS for the principal components

GWAS found 8 unique genes associated with the PCs characterizing the main modes of variation in TPCs (Table 3). No significant SNPs were found for the PC1 interpreted as a TPC elevation, even though it explained most of the phenotypic variation (Fig. 3A). However, the PC1 had a nearly significant association (P = 2.12 × 10^−8^) with the gene (NCU00170) encoding a DNAJ domain-containing protein, which is likely to be a true association based on its occurrence in the GWAS for growth rates at different temperatures (Table 2). The PC2 was interpreted as a shift in TPC optimum towards colder temperatures and it had 3 associations located in chromosomes 1 and 7 (Table 3, Fig. 6). The first association was upstream from the gene (NCU03028) encoding a deubiquitination-protection protein dph1, which works in protein degradation and localization to vacuoles, and has been found to affect mycelial growth under starvation stress (Zhou et al. 2016). The second association was in the gene (NCU06948) encoding an EF-hand domain-containing protein and had a very strong allelic effect (–0.99 mm/h). The EF-hand proteins affect enzymatic activity and cell division through microtubule organization and have been found to be involved in vesicular trafficking and fungal hyphal growth (Kundu et al. 2022). The third association was a missense variant having a large allelic effect (–0.85 mm/h) and changing alanine to serine in the *pks-4* (NCU08399) gene. The *pks-4* encodes a polyketide synthase 4 variant which functions in the formation of fatty acids. The polyketide synthases have been linked to the sexual reproduction and synthesis of secondary metabolites against predation, which both are important to the survival of *N. crassa* (Zhao et al. 2017).

**Table 3.** SNPs that were significantly associated with the predictive values of the first 3 principal components, their positions in genome (bp), allelic effects on phenotypic variance, allelic variant types and annotated proteins. The signs of the allelic effects are with respect to the minor allele and the annotations for intergenic variants are presented for the closest gene.

| PC | Chromosome | Position (bp) | P-value | Allelic effect | Gene | Symbol | Annotation | Variant |
| --- | --- | --- | --- | --- | --- | --- | --- | --- |
| 2 | 1 | 8 395 566 | $5.16 \times 10^{-9}$ | -0.14 | NCU03028 | | deubiquitination-protection protein dph1 | upstream |
| 2 | 7 | 871 838 | $4.21 \times 10^{-11}$ | -0.99 | NCU06948 | | EF-hand domain-containing protein | 3'UTR |
| 2 | 7 | 1 091 058 | $5.03 \times 10^{-9}$ | -0.85 | NCU08399 | *pks-4* | polyketide synthase 4, variant | missense |
| 3 | 1 | 7 807 362 | $6.30 \times 10^{-20}$ | 0.28 | NCU00621 | | TORC1 growth control complex subunit Kog1 | 5'UTR |
| 3 | 2 | 683 209 | $2.70 \times 10^{-10}$ | 0.12 | NCU03448 | | hypothetical protein | upstream |
| 3 | 2 | 3 468 329 | $7.39 \times 10^{-12}$ | -0.09 | NCU08496 | | hypothetical protein | missense |
| 3 | 3 | 337 373 | $2.91 \times 10^{-9}$ | -0.14 | NCU10002 | | hypothetical protein | synonymous |
| 3 | 5 | 4 421 459 | $4.56 \times 10^{-16}$ | -0.21 | NCU06730 | | hypothetical protein | upstream |

**Figure 6.**
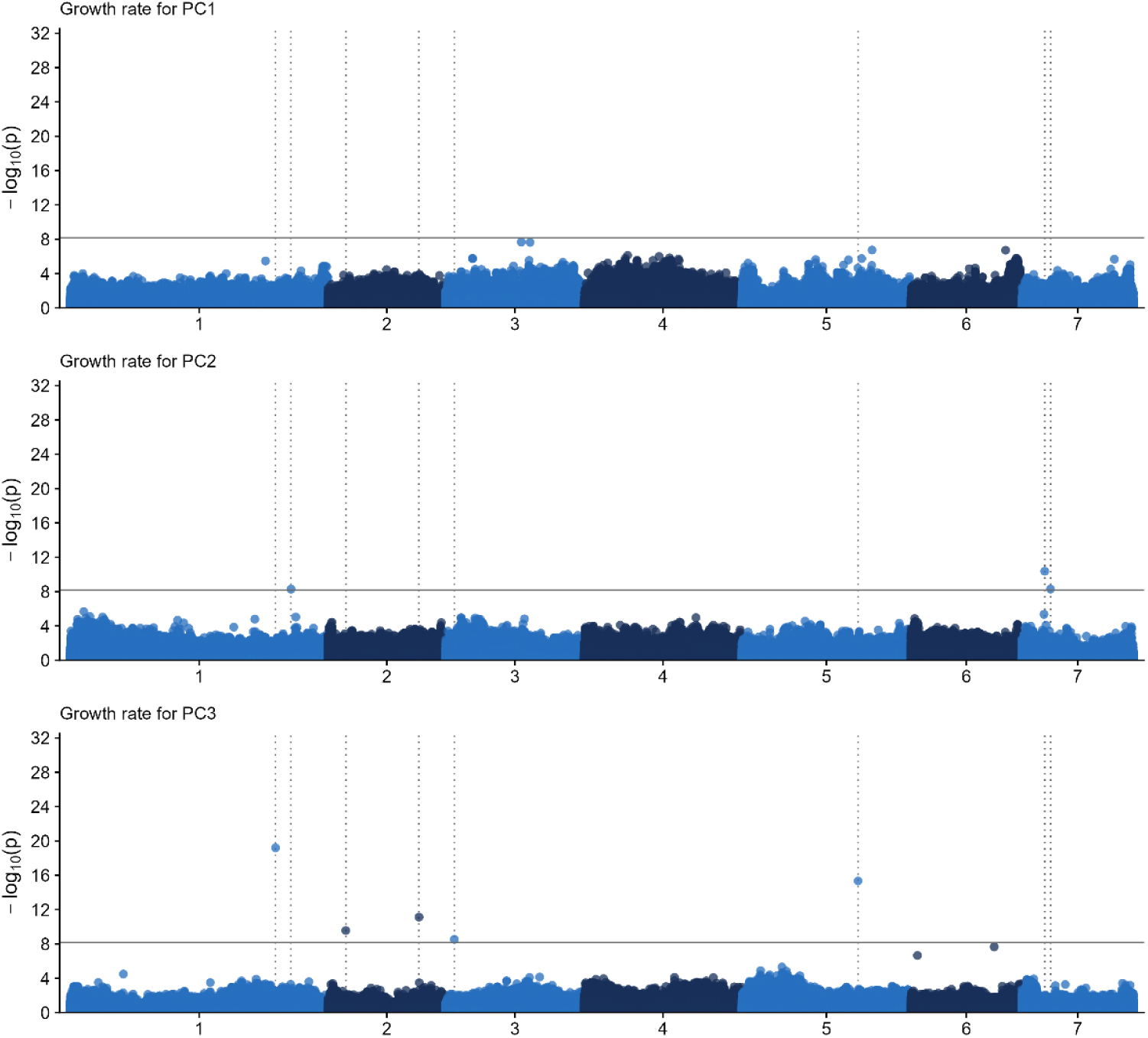
Manhattan plots for the variance in growth rate explained by the PC1, PC2 and PC3. The solid horizontal lines denote a P-value threshold of 0.01 after a Bonferroni correction. The dotted vertical lines highlight the candidate SNPs that were significant at least for one of the PCs.

The PC3 was interpreted as the shape change towards higher temperatures and had 5 associations in chromosomes 1, 2, 3, and 5 (Table 3, Fig. 6). The strongest association (P = 6.30 × 10^−20^) was in the gene (NCU00621) encoding a TORC1 growth control complex subunit Kog1, that was also associated at 42 °C, but with the PC3 there was a moderate positive allelic effect (Table 2, Table 3). There was also another association with positive allelic effect upstream from the gene (NCU03448) with unknown function (Table 3). In addition, 3 genes having negative allelic effects and encoding hypothetical proteins were associated with the PC3. One of them (NCU08496) was a missense variant changing alanine to serine and had a high minor allele frequency (0.45) in the mapping population. Other mutations were synonymous (NCU10002) and intergenic upstream (NCU06730) variants. In the previous GWAS for the growth rate of *N. crassa*, different genes were found to be important at optimal temperature, and at hot and cold extremes of the TPC (Räsänen et al. 2024).

## 4 DISCUSSION

Climate change obscures the increases in the mean and variance of temperature, yet it is not well understood how different thermal environments are selecting individuals, genotypes and genes (Kingsolver et al. 2013; Sinclair et al. 2016; Kristensen et al. 2020). A more thorough understanding of the genetic basis of thermal tolerance can be achieved with studies integrating quantitative and molecular genetics methods (Cortés et al. 2020; Buckley and Kingsolver 2021). To establish whether distinct genetic architectures are regulating the growth rate of *N. crassa* at constant and fluctuating temperatures, we estimated the amounts of genetic variation and covariation, and interpreted the directions of most evolutionary potential in TPCs. Interestingly, GWAS found many unique loci associated with growth rate either at constant or fluctuating temperatures, or with the PCs reflecting the modes of the TPCs. However, the general view of our results indicated a high level of shared genetic variance across temperatures which was due to the elevated overall performance.

We found that the constant extreme temperature 42 °C and fast 32–42 °C fluctuations had the most negative effects on the growth rate of *N. crassa*. Previous studies have suggested that the fast frequency of fluctuations might be more important to the organism’s fitness than the duration or the magnitude of the exposure to extreme temperatures (Kearney et al. 2012; Marshall and Sinclair 2015). In *N. crassa*, the upregulation of the heat shock protein (HSP) production is activated in minutes at temperatures around 42 °C (Mohsenzadeh et al. 1998; Kapoor and Roy 2014), but the HSP synthesis requires energy, which generally leads to extensive fitness costs after repeated exposures to stressfully high temperatures (Sinclair et al. 2016). In a cell, the allocation of energy to stress-related functions are known to mutually repress growth-related pathways such as protein synthesis and ribosome biogenesis (López-Maury et al. 2008). This mechanism is called a ‘general stress response’, which redirects the resources from growth to stress resistance as a response to multiple stresses, meaning that the most stress-resistant cells are non-growing (Kültz 2005). Under faster fluctuations, there is also less time to acclimate. In *N. crassa*, there is a peak in HSP production in around 60 min (at fast fluctuations, one step lasts 15 min), after which the recovery period with declining mRNA levels and HSP degradation starts (Kapoor and Roy 2014). It has been found that when temperature fluctuations became faster than the recovery time of *N. crassa*, the growth rate slows down also at more optimal temperatures, as the recovery starts to take longer than the acclimation to heat exposure (Kronholm and Ketola 2018).

The quantitative genetics analyses gave us estimates of the genetic variation in several small-effect loci. We found high heritability values across temperatures, meaning that most of the variation in growth was due to genetic variation among the strains. We found also strong genetic correlations between the fast and slow frequency fluctuations, and between fluctuations and their mean temperatures, predicting correlated responses to the selection (Czesak et al. 2006). This indicated that in *N. crassa*, there should not be strong trade-offs or constrains between adaptation to constant and fluctuating temperatures. However, the genetic correlations and covariances with the high extreme temperature 42 °C were markedly lower, pointing that the growth at 42 °C could evolve more independently from the selection on fitness at other temperatures. Similar quantitative genetic results have been found before for the growth rate of *N. crassa* at constant 40 °C (Moghadam et al. 2020), suggesting that different genes or amounts of gene expression might affect acclimatization at the higher thermal limit.

The high heritability values and strong positive genetic correlations between environments might be found because some of the strains are superior in growth rate, so called ‘supergeneralists’ or ‘master of all temperatures’ (Huey and Hertz 1984; Kristensen et al. 2020). We verified this suggestion by PCA and found that most of the variation (PC1, 83 %) was in the individual’s overall fitness in growth rate regardless of the temperature, meaning that some strains grew better, and some slower across all the tested temperatures. Previously, the TPC elevation has been found to predominate the growth rate in *N. crassa* (Moghadam et al. 2020) and performance in other species (Yamahira et al. 2007; Shama et al. 2011; Ketola et al. 2014; Latimer et al. 2015; Bartheld et al. 2017; Schaum et al. 2022). These observations are on the contrary to the traditional hypothesis of specialist-generalist trade-offs preventing the evolution of superior generalists (Angilletta 2009), and some studies that have found costs between performances at intermediate and extreme temperatures (reviewed in Logan and Cox 2020). According to a hypothesis of condition-dependent traits, the fact that many studies have failed to find evidence for the costs of plasticity (Berger et al. 2014; Latimer et al. 2015; Murren et al. 2015; Schaum et al. 2022), can be explained by a larger amount of genetic variation in individual’s overall fitness traits compared to stress tolerance (Kristensen et al. 2020; Rowe and Houle 1996; Van Noordwijk and De Jong 1986). Based on this, the superiority in fitness could arise due to more efficient resource acquisition, metabolic assimilation, and allocation on dependent traits across temperatures. Indeed, it has been found that the traits associated with individual’s overall fitness are usually those related to cells resource uptake and energy efficiency (Kristensen et al. 2005; Pedersen et al. 2008). In addition to allocation of energy in growth across temperatures, allocation trade-offs can exist between correlated traits sharing the same functional or developmental basis (Angilletta et al. 2003; Berger et al. 2014).

We found much less variation explained by the subsequent PCs, indicating little evolutionary potential in TPCs for the optimum shift towards colder temperatures (PC2, 8 %) and for the shape change to better performance at high temperatures (PC3, 4 %). The loadings of the PC2 were interesting since the tolerance to constant temperatures seemed to also affect the tolerance to fluctuations within the same temperature range, indicating that in our mapping population, there were, to some extent, cool and warm adapted genotypes (Angilletta 2009; Berger et al. 2014). The PC3 showed little genetic variance in tolerance to high extreme temperatures, which most likely means the induction or the magnitude of the heat-shock response (Moghadam et al. 2020). One possibility is that our nature-derived strains have undergone selection during their thermal history and are already well-adapted to heat, decreasing the amount of variation in this tolerance trait. The ability to tolerate prolonged heat is known to be a trait that is highly evolutionary conserved as temperature affects all biochemical reactions (Araújo et al. 2013; Hochachka and Somero 2002). Many experiments have found little variation in genomic regions important to heat stress, suggesting evolutionary constraints on heat tolerance (Mitchell and Hoffmann 2010; Kellermann et al. 2012; Kristensen et al. 2015). However, some studies have found evidence that species can expand their heat tolerance if there is a strong selection for better performance at hot temperatures, which is often observed in evolution and selection experiments using constantly high temperatures (Bennett et al. 1990; Holder and Bull 2001; Bubliy and Loeschcke 2005; Hangartner and Hoffmann 2016).

By GWAS we discovered unique large-effect loci associated with growth rate only at constant temperatures or fluctuations of both ranges and frequencies, but also genes that had shared associations at constant and fluctuating temperatures. Interestingly, at temperatures close to extremes only unique SNPs were associated, whereas shared SNPs were associated at more optimal temperatures, pointing that adaptation to extremes could be more specific. Also, there were no shared SNPs between the fast and slow fluctuations. Most of the allelic variants did not affect the protein structure or were at the regulatory area of the genes, being potential regulatory variants. However, the types of allelic variants were not associated clearly with growth at constant or fluctuating temperatures, even though regulatory area mutations are often expected to dictate the dynamic responses under thermal fluctuations (López-Maury et al. 2008). Almost all minor alleles slowed down the growth rate and no opposite allelic effects on growth were found at different temperatures for the shared SNPs. This supported our findings from quantitative genetics, which didn’t show strong evidence for the existence of trade-offs. The predominantly negative allelic effects and the absence of any clear trade-offs have been found also in previous studies on thermal tolerance in *N. crassa* (Moghadam et al. 2020; Räsänen et al. 2024).

There was not much overlap in SNPs associated with growth rate at different temperatures and with the PCs, but the overlapping molecular functions of the associated genes suggested a shared physiology, probably owing to the general stress response (López-Maury et al. 2008; Gerken et al. 2015). Our genome-wide analysis found candidate genes that encoded proteins essential for the basic functions in fungi, such as transcription, stress resistance, cellular growth, and metabolism. Many of these proteins are known to be relevant to the ecology and fitness of *N. crassa* and showed sensitivity to temperature or other environmental stresses. It has been found that at lower temperatures, genes having preparatory functions in thermal tolerance are activated, for example, preventing the damage on cell structure through membrane composition and anticipatory production of stress proteins (Lecheta et al. 2020). At higher temperatures more functions should be in dynamic stress protein production, for example, in the upregulation of the HSP synthesis to repair the unfolded proteins (Sørensen et al. 2003; Lecheta et al. 2020). Our results were in line with these assumptions, as at temperatures up to optimum, proteins functioned in transcription, ribosome biogenesis, nutrient uptake, and energy production. At temperatures of both lower and higher range, and especially at fast fluctuations, we found a regulatory area mutation in DNAJ (HSP40) protein, and there was also a nearly significant association with the TPC elevation. At constant 37 °C the encoded proteins worked in signal transduction and there was already heat related regulation for growth. At higher range 32–42 °C fluctuations we found genes that affected the folding and thermodynamic stability of the proteins. It has been suggested that at stressfully high temperatures, the adaptation can be driven also by other genes than HSPs because of the costs of maintaining a dynamic stress response (Sørensen et al. 2003). At constant 42 °C and with PC3 we found TORC1 that regulates mycelial growth under heat stress (Urban et al. 2007; López-Maury et al. 2008), supported by the negligible growth rates observed at high extreme temperature.

Overall, our results indicated that there is a shared polygenic basis of thermal tolerance in *N. crassa* at constant and fluctuating temperatures. This tolerance seems to evolve mainly via elevated TPCs, which could be due to individual’s overall fitness or the general stress response. Therefore, the increasing variance in temperature should not set bounds on the evolutionary potential of *N. crassa*, but the ability to adapt to extreme heat is probably more constrained. The identified candidate genes are also likely to be important for the thermal tolerance in growth rate, which is an important fitness trait related to both pathogenicity and mutualism in filamentous fungi (Pringle and Taylor 2002). As climate change keeps escalating, more research is needed to scrutinize the genetic basis of thermal tolerance and the species ability to adapt to various fluctuations. To this end, we argue that quantitative genetics and molecular methods like GWAS should be used together, since solely these methods do not provide a complete picture of the genetic architecture and might result in opposite interpretations.

## Acknowledgements

The authors wish to acknowledge CSC – IT Center for Science, Finland, for computational resources and the Fungal Genetics Stock Center for the strains used in this study. The best thanks also to Neda Moghadam and Karendeep Sidhu for their part in the data collection. This study was funded by the University of Jyväskylä Doctoral Programme in Biological and Environmental Science (ER), and grants from Emil Aaltonen foundation (ER), and the Academy of Finland (TK: 278751, IK: 274769 and 321584).

## Author contributions

IK, TK, and ER planned the questions and the experiment together. ER, ML, PAMS and IK conducted the laboratory work. IK and ER run the GWAS and other statistical analyses with advice from TK. ER was responsible for writing the manuscript, and all authors have read and approved the current version. Authors declare no conflicts of interests.

## Data access

Genotypic data for the mapping population can be found at: XXXX.doi, and the phenotypic data and scripts at YYYY.doi.

